# Isothermal Detection of Influenza D using RT-LAMP

**DOI:** 10.64898/2026.01.09.698603

**Authors:** Carlos Abelardo dos Santos, Jialu Li, Pauline M. van Diemen, Andrew McMahon, Benjamin C. Mollett, Andrew M. Ramsay, Meshach M. Maina, Joe James, Helen E. Everett, Janet M. Daly, Nicole C. Robb

## Abstract

The *Orthomyxoviridae* family includes influenza D virus (IDV), an emerging pathogen primarily affecting cattle and swine, with evidence of cross-species transmission and potential zoonotic risk. Although active human infections have yet to been confirmed, high seroprevalence in cattle-exposed populations highlights the need for continued surveillance. Here, a rapid, field-deployable RT-LAMP assay for IDV detection was developed and validated, with 99.2% specificity and sensitivity ranging from 95.6% (Cq < 30) to 81.8% (Cq < 40). This method offers a cost-effective, accessible alternative to RT-qPCR, enabling improved monitoring of IDV, and reinforcing preparedness for emerging influenza threats.

## Introduction

The family *Orthomyxoviridae* comprises nine genera, four of which are influenza viruses. Influenza A (IAV) and influenza B (IBV) are the main viruses responsible for human seasonal influenza epidemics, whilst IAV poses a pandemic threat linked to bidirectional transmission between animals to humans. Both IAV and IBV can cause severe illness in humans, whilst influenza C virus (ICV) infection is associated with milder upper respiratory tract symptoms, most commonly in children under the age of 2. Influenza D virus (IDV) was first discovered in 2011 in pigs in Oklahoma, USA, and subsequently found in cattle, which are now considered the main livestock reservoir. In cattle, IDV infection is associated with bovine respiratory disease (BRD) as the primary viral infection, which may predispose cattle to opportunistic secondary bacterial infections (1).

Whilst the majority of IDV occurrences have been in swine and cattle, IDV antibodies have also been found in other domesticated animals such as sheep (2), goats (3), horses (3), wild boars (4) and camelids (5), suggesting that cross-species transmission may occur. Mice, ferrets and guinea pigs have also been found to be susceptible to experimental viral infection (6). Similarly to ICV, IDV has been shown to bind to sialic acid receptors on the cell surface of the host, specifically 9-O-acetylated sialic acid receptors (7). These receptors are found throughout the entire respiratory tract in cattle, as well as in the nasal and pharyngeal epithelium of pigs, sheep, goats, and horses, suggestive of a potentially wide host range (8).

Influenza D virus has been shown to propagate effectively in various human cell types (9), however, no active infection of humans has been reported to date, and although IDV was detected in a nasal wash sample from a swine farm worker in Malaysia, no infectious virions were retrieved (10). The presence of IDV antibodies has been the primary method of inferring past exposure to IDV in humans. A 2011 study found a 1.3% seroprevalence of IDV antibodies in the general human population in the USA and Canada (11). Another study in Italy discovered that the seroprevalence of IDV in the general human population increased from 5% in 2005 to 46% in 2014 (12). In human cohorts occupationally exposed to cattle, the prevalence of IDV antibodies reached as high as 97%(13), reflecting a similar prevalence of antibodies in cattle (14).

Although IDV infection has not been observed to induce severe disease in humans, it remains crucial to monitor its prevalence due to the ability of influenza viruses to evolve and transmit across species barriers. While antibody detection serves as a robust epidemiological tool on a population scale (15), reverse transcriptase quantitative polymerase chain reaction (RT-qPCR) stands as the gold standard for detecting viral RNA and thus active infection at an individual level. However, the necessity for specialized laboratories and trained personnel to process and analyse the samples, coupled with high costs and limited utility for field-based testing, poses a significant challenge to high throughput. Reverse transcription loop-mediated isothermal amplification (RT-LAMP) is a rapid, highly sensitive, and cost-effective molecular diagnostic tool that has emerged as a potential alternative to PCR-based methods (16). RT-LAMP can be performed at a constant temperature and return a result in less than an hour, making it well suited for point-of-sample testing. The success of RT-LAMP during the COVID-19 pandemic underscores its potential as a valuable tool in our ongoing efforts to combat infectious diseases (17).

Here, we report an RT-LAMP assay to specifically detect IDV with particular application for field setting diagnostic assays in resource-poor settings. We tested a total of 28 spiked samples and 46 isolated RNA samples from swine and cattle using the assay, which achieved a specificity of 99.23% and a sensitivity ranging from 95.60% for samples with a Cycle quantification (Cq) < 30, to 81.82% for all samples up to a Cq of 40. This development not only enhances our ability to detect and monitor IDV but also highlights the potential of RT-LAMP as a valuable tool in monitoring infectious diseases. We believe that our findings will support ongoing efforts to mitigate the risks associated with this virus and enhance our collective capacity to respond to infectious diseases.

## Materials & Methods

### Virus Isolates

The isolates D/swine/Oklahoma/1334/2011 (11) and D/bovine/France/5920/2014 (18,19) were propagated in embryonated chicken eggs and on hRT18G cells respectively. Both strains are well characterized and produced with a minimum number of passages to avoid loss of strain fidelity. D/swine/England/126471/2023 is a more recent isolate and was propagated on ST cells (20).

### RNA extraction

We carried out the pre-isolation steps depending on the nature of the sample. For viral isolates, 20 μL of the cell supernatant was lysed with 350 μL of Monarch StabiLyse DNA/RNA Buffer, and we carried out the extraction according to the manufacturer’s protocol. We also extracted RNA from a swine lung fragment by excising a 10 mg tissue fragment and lysing it in 200 μL of StabiLyse DNA/RNA Buffer and 200 μL of nuclease-free water (Integrated DNA Technologies) in a FastPrep-24 bead beater homogenizer (MP Biomedicals) for approximately 10 minutes. We extracted total RNA from each preparation using the Monarch Spin RNA Isolation Kit (New England Biolabs), according to the manufacturer’s instructions.

### RT-qPCR assay for IDV

The RT-qPCR primers and probes for IDV detection used in this study were originally published by Faccini et al. (21). The reactions were performed in a 384-well Quant Studio 5 Real-Time PCR System (Applied Biosystems) using the Luna Universal Probe One-Step RT-qPCR Kit (New England Biolabs) following the manufacturer’s instructions. The primers and probe target the PB1 gene (from position 1215 to 1323, Accession number JQ922306), so we designed a PB1 gBlock to use as control containing the nucleotides from position 500 to 1500. The gBlock was synthesized by Integrated DNA Technologies. We eluted the gBlock according to the manufacturer’s protocol and confirmed the concentration by UV absorbance using a NanoDrop spectrophotometer (Thermofisher Scientific). We used serial dilutions ranging from 10^6 to 10 copies of the IDV gBlock to generate a standard curve, which was employed to quantify the RNA material from isolates, mock samples and samples.

### RT-LAMP Primer Design

We downloaded all the PB1 and PB2 complete gene sequences of IDV available from the National Centre for Biotechnology Information (NCBI) database (accessed December 2024) and generated alignments using the MAFFT multiple sequence alignment tool (https://mafft.cbrc.jp/alignment/software/). We designed the primers for RT-LAMP against PB1 and PB2 using the PrimerExplorer V5 software (https://primerexplorer.eiken.co.jp/lampv5e/index.html). To improve coverage and guarantee amplification of the maximum number of sequences, we set the consensus threshold to 90%. We marked all the mutated positions in the primer design software and created the primers using the “Common” design option, which allows for primers to be designed targeting mutated regions if the position of the variable nucleotide within the primer is unlikely to affect amplification efficiency. Primers were filtered by end stability, and the maximum ΔG for the 3’ end of F3/B3, F2/B2, LF/LB and the maximum ΔG for the 5’ end of F1c/B1c was set to be ≤ -4.00 kcal/mol. Primers were also checked for the presence of stable hairpins, self-dimers and homodimers using IDT’s Oligo Analyzer tool (https://eu.idtdna.com/pages/tools/oligoanalyzer). Sets containing primers with stable hairpins or dimers were not synthesised. The primers FIP and BIP are composed of two different regions (F1c/F2 for FIP and B1c/B2 for BIP), which mediate the creation of the loops characteristic of LAMP. Target recognition by LAMP primers creates an ever-growing concatemer containing single-stranded loops that also serves as target for continuous amplification. To decrease local Tm in the loops and lower the chances of undesired secondary structures that could hinder amplification, we inserted T linkers (22) between the F1c/F2 and B1c/B2 regions in the FIP/BIP primers.

### Colorimetric one-step RT-LAMP assay for IDV

We prepared all reactions as a total volume of 20 μL, each containing a final concentration of 1X WarmStart® Colorimetric LAMP Master Mix with UDG (Uracil DNA Glycosylate) (New England Biolabs), 1X primer mix (containing 0.2 mM F3/B3, 1.6 mM FIP/BIP and 0.8 mM LF/LB primers), 10% (v/v) of target, and nuclease-free water (Integrated DNA Technologies). For real-time analysis, we added 1 mM of SYTO9 (Thermofisher Scientific) to the reaction mixture. To optimize the assay, we incubated the reactions in a real-time thermocycler (QuantStudio 3, Applied Biosystems) at 65°C for 80 cycles, each cycle consisting of 45 seconds of incubation followed by fluorescence read in the FAM channel for 15 seconds. Subsequent reactions were incubated in a conventional T-100 PCR instrument (Bio-Rad) at 65°C for 60 minutes. Unless otherwise specified, all reactions were performed in triplicate.

### Sensitivity and specificity of the Influenza D assay

To assess whether we could differentiate IDV from closely-related viruses, we tested the RT-LAMP assay for IDV against gBlocks containing the PB2 target gene segment of all other influenza strains. While the gBlock containing the complete PB2 gene for IDV (D/swine/Oklahoma/1334/2011) was used as positive control in our reactions, gBlocks containing the complete PB2 gene for Influenza A (IAV, A/Puerto Rico/8/1934), Influenza B (IBV, B/Lee/1940), and Influenza C (ICV, C/Ann Arbor/1/50) viruses were used as negative controls. We carried out the reactions using either 2 μL of the respective gBlocks controls or the equivalent amount of nuclease-free water as a non-template control (NTC). All the gBlocks were tested at a concentration of 10^6^ copies per μL for 1 hour. Results were analysed in triplicate, based on the colorimetric change in the reaction from pink (negative) to yellow (positive).

To determine the sensitivity of the assay, we mixed equimolar amounts of extracted RNA from each of the three viral isolates (D/bovine/France/5920/2014, D/swine/Oklahoma/1334/2011 and D/swine/England/126471/2023), which had been quantified by RT-qPCR using the assay described previously. We serially-diluted the RNA mix and verified the concentrations by RT-qPCR. We tested the serially diluted RNA mix by RT-LAMP in octuplicate and fitted a Probit regression curve (dose-response plot) using the IBM SPSS Statistics Software (Version 29.0.2.0). We defined the limit of detection to be equal to the minimum number of copies detected in at least 95% of the reactions.

We analysed the amplification products on a 2% agarose gel in 1X TBE (Tris-Borate-EDTA) Buffer using SYBR Safe DNA Gel Stain (Thermofisher) and the Quick-Load 100 bp DNA Ladder (New England Biolabs). The gel was run at 100V for 60 minutes and images were acquired with a ChemiDoc Imaging System (BioRad).

### Evaluation using spiked and field-collected animal samples

To assess the performance of the test in a more complex sample matrix, we prepared 28 spiked samples by mixing total purified RNA extracted from swine lung tissue with variable amounts of viral RNA from the three virus isolates used in this study. Samples were quantified by RT-qPCR and tested by RT-LAMP. Results were analysed by the colour change from pink (negative) to yellow (positive) in the reaction after 60 minutes of incubation.

To assess the viability of the assay in real animal samples, we analysed 46 swine and cattle RNA samples. These samples comprised nasal swab and/or respiratory tissue samples (lung, viscera) from pigs and cattle with respiratory disease signs (cough and/or dyspnoea) submitted through the passive APHA surveillance networks, and containing a mixture of IAV or IDV positive or negative samples

The specificity and sensitivity of the RT-LAMP assay for detecting IDV was assessed by comparing its results with those of the RT-qPCR assay, using the free online tool Medcalc (available at https://www.medcalc.org/calc/diagnostic_test.php).

## Results

### Optimization of an RT-LAMP assay for Influenza D

The primer sets generated for the PB1 and PB2 gene sequences of IDV were evaluated in silico for their end stabilities, formation of hairpins, self-dimers and heterodimers. We found that none of the sets targeting the PB1 gene passed our filters, while two primer sets targeting the PB2 gene met our criteria and were synthetized. The sequences of the two RT-LAMP primer sets that passed our filters are presented in Table S1.

We initially trialled both primer sets by performing the RT-LAMP reaction in a real-time thermocycler targeting gBlocks containing the complete PB2 sequence of IDV, as well as IAV, IBV and ICV as negative controls, using SYTO9 DNA dye to track the accumulation of amplification products over time (Figure 1). Each gBlock was incubated at a concentration of 10^6^ copies per µL of input. As non-specific amplification is a well-known problem with RT-LAMP reactions, we incubated the reactions for up to 80 minutes to assess the maximum incubation time for each primer set, defined as the longest time in which reactions could proceed without producing false-positive signals in the negative samples or non-template controls. While both primer sets amplified the Influenza D target gBlock at around the 15 minute mark, the IDV_set1 showed early onset of non-specific amplification at around 40 minutes (Figure 1A), compared to 65 minutes with the IDV_set2 (Figure 1B). Due to the increased specificity, IDV_set2 was therefore chosen for all subsequent analysis.

**Figure 1.**
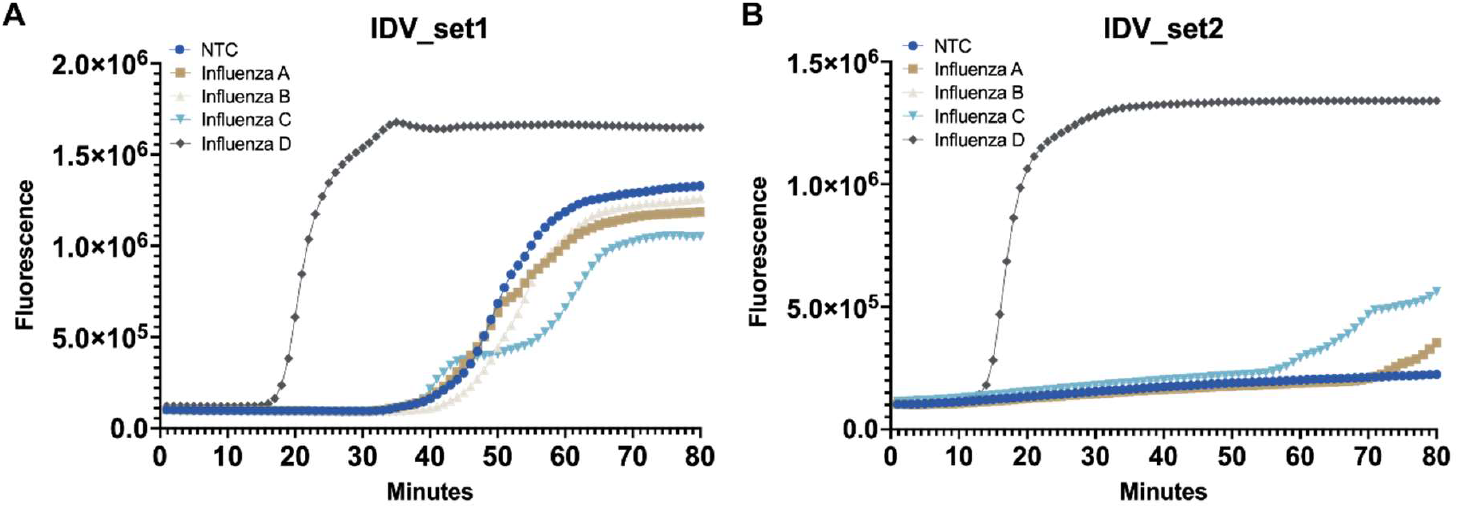
Mean amplification curves from RT-LAMP reaction triplicates showing the onset of specific and non-specific signal over time. RT-LAMP on gBlocks containing the influenza A, B, C and D virus PB2 gene sequences. All gBlocks were tested at a concentration of 10^6^ copies per µL of input. A) Reactions using the primers in IDV_set1. The amplification of the IDV gBlock positive control started within the first 17 minutes, whilst IAV, IBV and ICV gBlocks and the NTC produced false-positive signals after only 35 minutes of incubation. B) Reactions using the primers in IDV_set2. The amplification of the IDV gBlock positive control started within the first 15 minutes, whilst IAV, IBV and ICV gBlocks and the NTC produced false-positive signals after 65 minutes of incubation. NTC: Non-template control (water).

To ensure complete coverage of at least 90% of the published IDV lineages, including the three viral isolates that we had access to (D/bovine/France/5920/2014, D/swine/Oklahoma/1334/2011 and D/swine/England/126471/2023), degenerate nucleotides were incorporated into the primers from IDV_set2 where necessary (Table S1 and Figure 2A). To test whether the primer set containing degenerate nucleotides enabled detection of all three available isolates of IDV, we incubated dilutions of extracted RNA at 10^4^ and 10^3^ copies/µL for each of the isolates. All concentrations of the three isolates were efficiently amplified after approximately 15 minutes (Figure 2B).

**Figure 2.**
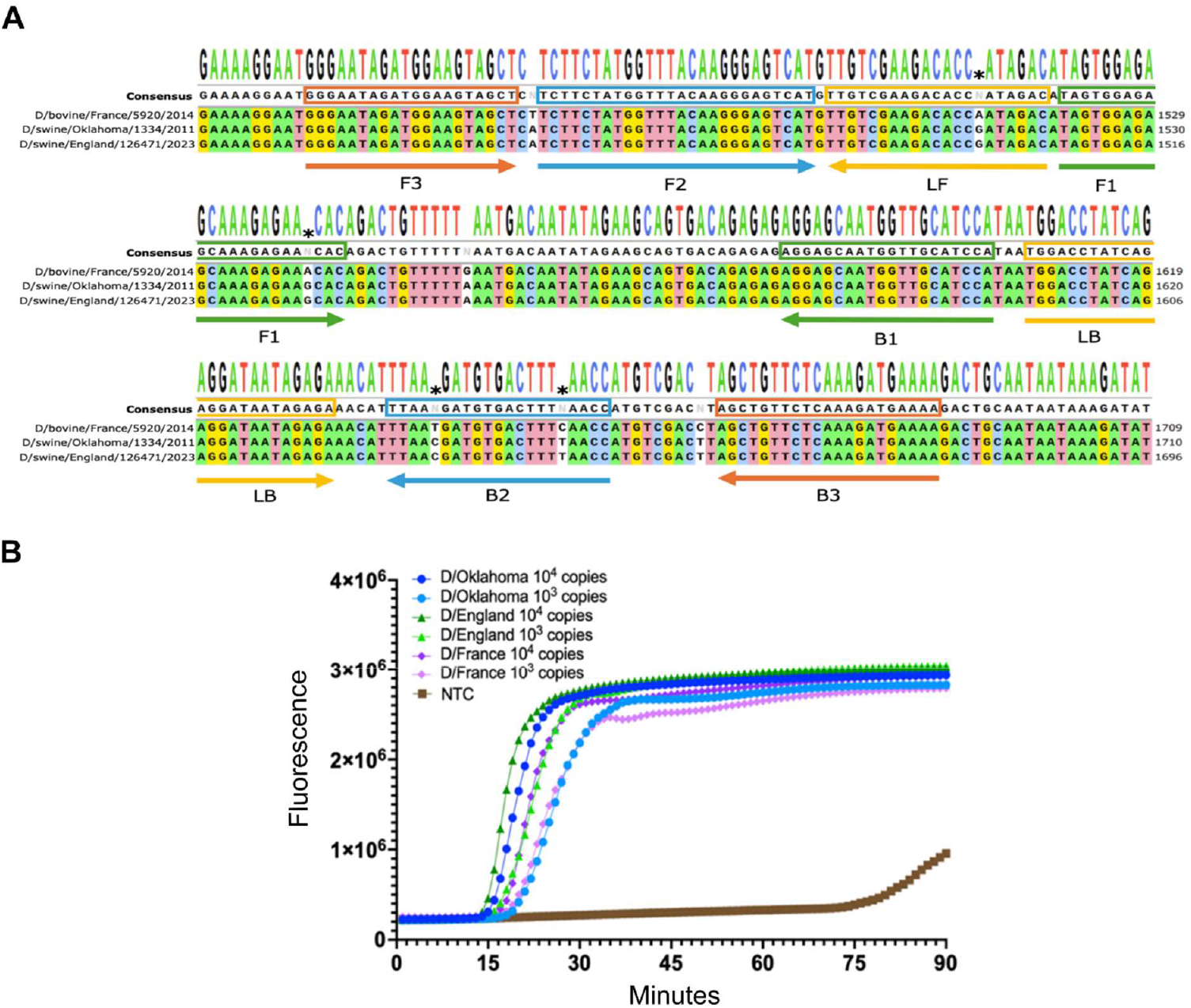
RT-LAMP primers targeting the influenza D virus (IDV) PB2 gene efficiently amplify viral isolate RNA. A) All available IDV PB2 sequences in the NCBI database were aligned and mutations were annotated. To improve coverage, primers were synthesised with degenerate nucleotides in positions with heterogeneity in the sequences. The consensus sequence and sequences of the three isolates, D/bovine/France/5920/2014, D/swine/Oklahoma/1334/2011, and D/swine/England/126471/2023 are shown for comparison. Primer target sequences are presented within the boxes, and the direction of binding is represented by the arrows. B) Mean amplification curves from RT-LAMP reaction triplicates showing the onset of specific and non-specific signal over time. All IDV isolate RNAs at 10^4^ and 10^3^ copies per µL amplified within the first 20 minutes regardless of the lineage. NTC: Non-template control (water).

### The colorimetric RT-LAMP assay exhibits high sensitivity and specificity

Having shown that the RT-LAMP primers efficiently detected IDV RNA using a real-time thermocycler, we incubated subsequent reactions in a conventional thermocycler and analysed the results visually based on a colorimetric change in the reaction from pink (negative) to yellow (positive).

To show that the addition of the degenerate nucleotides did not impact primer specificity, a high number of gBlock copies (10^6^ copies/µL) of the PB2 gene of influenza A, B, C and D were incubated with the RT-LAMP mixture for 60 minutes at 65°C, only the IDV target was amplified (Figure 3A), suggesting that our assay is specific to IDV. This result was confirmed via gel electrophoresis of the amplicons (Figure 3B).

**Figure 3.**
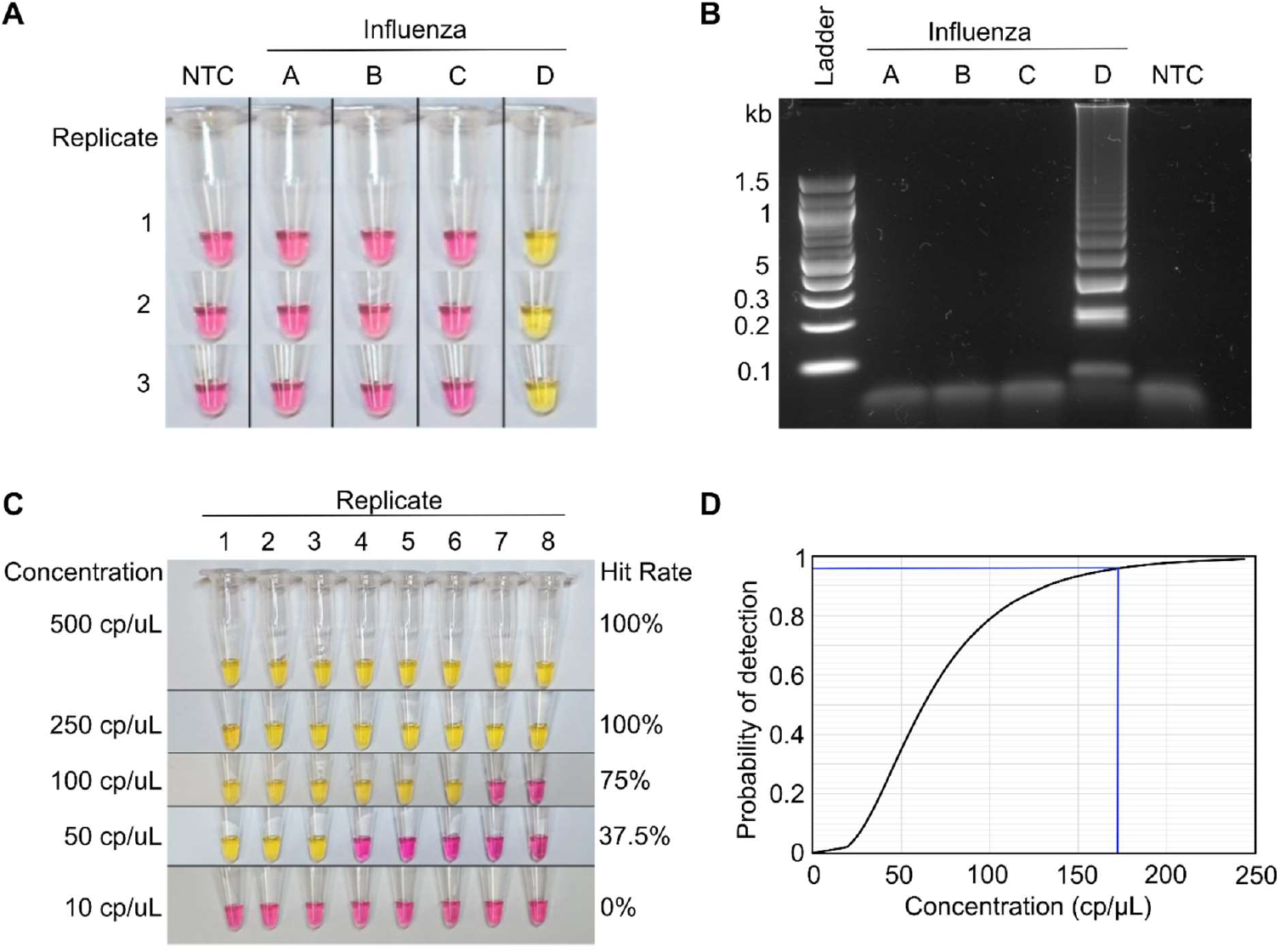
Specificity and sensitivity assays for influenza D virus (IDV) detection using RT-LAMP. A) The total of 10^6^ copies/µL of gBlocks containing the influenza A, B, C and D virus PB2 gene sequences were tested. Only the gBlocks containing the target sequences were amplified as seen by the colour change in the reactions. NTC: non-template control (water). B) 2% gel electrophoresis of one representative of each amplification products of the RT-LAMP reactions shown in A), showing the typical lamp ladder-like pattern in IDV virus samples and no visible non-specific bands in the other target samples. C) Dilutions of target RNA in eight replicates for sensitivity analysis at different concentrations, varying from 500 copies per µL (cp/μl) of target (top row) to 10 copies per µL of target (bottom row). D) The probit-analysis curve of probability of detection in relation to the number of copies. The limit of detection is defined as the minimum number of copies detected >95% of the time.

Next, the sensitivity of the essay was estimated from the amplification results of eight replicates at low target concentrations (500, 250, 100, 50 and 10 copies per µL of input) using an equimolar mix containing the RNA from the three IDV isolates used in this study. The calculated probit regression curve was plotted and the minimum number of copies detected 95% of the time was estimated to be 167 copies/µL (95% C.I. 103–3115 copies/µL) (Figure 3C&D).

### Evaluation of RT-LAMP assay on spiked and diagnostic samples

To assess the efficiency of the RT-LAMP test in a more complex matrix, we spiked varying concentrations of IDV RNA into total RNA extracted from swine lung tissue. Prior to addition of the IDV RNA the swine lung tissue was confirmed to be free of IDV RNA via RT-qPCR. Twenty-seven spiked samples were tested in duplicate in different amounts of swine lung RNA to mimic different target/background ratios, using 4 non-spiked samples as negative control (a total of 58 individual RT-LAMP reactions). The IDV RNA in all spiked samples was also quantified by RT-qPCR. The RT-LAMP assay successfully detected all samples that had a Cycle quantification (Cq) number lower than 29. It failed to amplify 2 samples out of 16 with a Cq between 29 and 30, 4 samples out of 14 with a Cq between 30 and 32 and 3 samples out of 4 with a Cq over 32 (Figure 4A). All uninfected swine lung RNA samples were negative.

**Figure 4.**
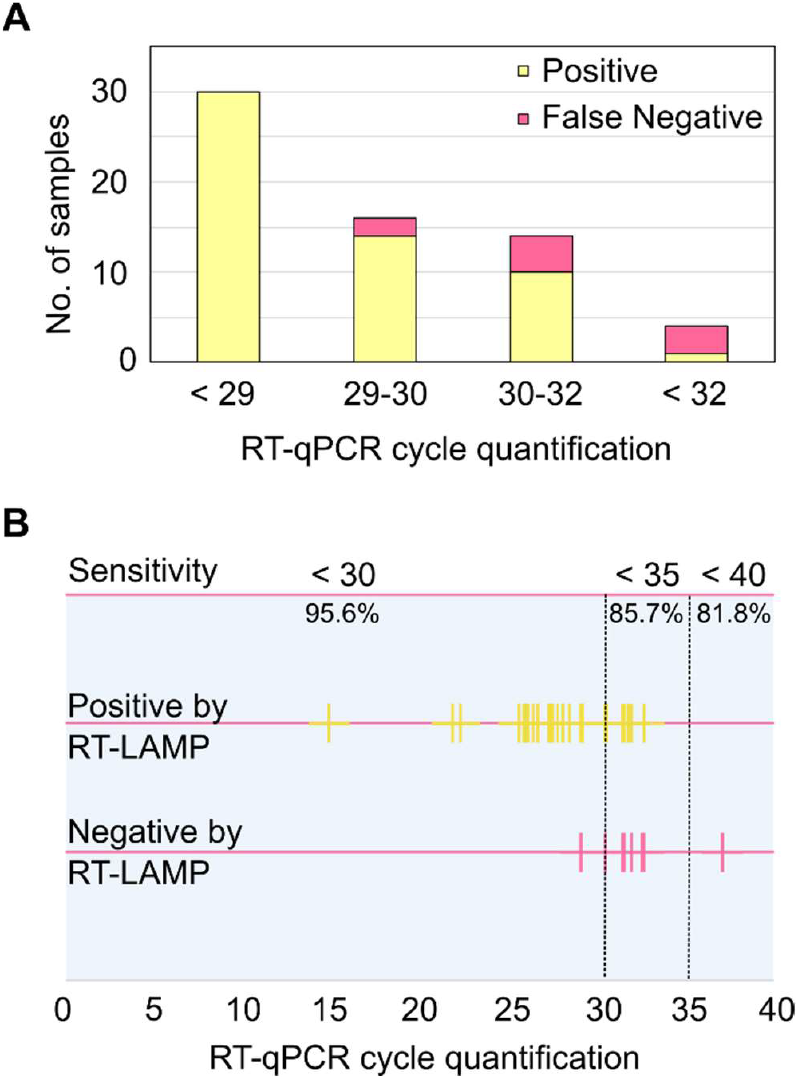
Sensitivity of the test in relation to RT-qPCR cycle quantification. A) Detection of spiked samples detected by RT-LAMP in relation to the cycle quantification (Cq) value. B) Each true positive sample was sorted according to its Cq value and RT-LAMP result. Yellow lines presented along the top line are positive by both RT-LAMP and RT-qPCR, pink lines on the lower line are positive by RT-qPCR but negative by RT-LAMP. The sensitivity of each range is presented as percentages at the top of the graph.

Finally, we tested the performance of the RT-LAMP assay in detecting IDV viral RNA in total RNA extracted from samples collected from animals in the field. We analysed a total of 46 RNA samples extracted from cattle and swine. The samples were composed of negative samples, samples containing IDV and samples containing IAV RNA. They were tested in triplicate and the results compared to RT-qPCR as shown in Table 1. The RT-LAMP assay successfully detected IDV RNA in 3 out of the 4 animal samples that were positive for IDV RNA by RT-qPCR.

**Table 1.** Comparison between RT-qPCR and RT-LAMP results for animal RNA samples. IDV: Influenza D Virus, IAV: Influenza A Virus. Samples 1-32 are from swine and 33-46 are from cattle. * Sample number 40 showed a change in colour in one of the replicates that did not fully convert into yellow, but for statistical calculations it was considered a false positive.

| Sample | IDV<br>RT-LAMP | IDV<br>RT-qPCR | IAV<br>RT-qPCR |
| --- | --- | --- | --- |
| 1 | Negative | Negative | Negative |
| 2 | Negative | Negative | Positive – Cq 36.45 |
| 3 | Negative | Negative | Negative |
| 4 | Negative | Negative | Negative |
| 5 | Negative | Negative | Negative |
| 6 | Negative | Negative | Negative |
| 7 | Negative | Negative | Negative |
| 8 | Negative | Negative | Positive – Cq 39.24 |
| 9 | Negative | Negative | Negative |
| 10 | Negative | Negative | Negative |
| 11 | Negative | Negative | Negative |
| 12 | Negative | Negative | Negative |
| 13 | Negative | Negative | Negative |
| 14 | Negative | Negative | Positive – Cq 32.19 |
| 15 | Negative | Negative | Negative |
| 16 | Positive | Positive – Cq 22.29 | Negative |
| 17 | Negative | Negative | Negative |
| 18 | Negative | Negative | Negative |
| 19 | Negative | Negative | Negative |
| 20 | Negative | Negative | Positive – Cq 27.88 |
| 21 | Negative | Negative | Negative |
| 22 | Negative | Negative | Negative |
| 23 | Negative | Negative | Negative |
| 24 | Negative | Positive – Cq 37.09 | Negative |
| 25 | Negative | Negative | Negative |
| 26 | Negative | Negative | Negative |
| 27 | Negative | Negative | Negative |
| 28 | Negative | Negative | Negative |
| 29 | Negative | Negative | Negative |
| 30 | Negative | Negative | Negative |
| 31 | Negative | Negative | Negative |
| 32 | Negative | Negative | Negative |
| 33 | Negative | Negative | Negative |
| 34 | Negative | Negative | Negative |
| 35 | Positive | Positive – Cq 14.84 | Negative |
| 36 | Negative | Negative | Negative |
| 37 | Negative | Negative | Negative |
| 38 | Positive | Positive – Cq 21.81 | Negative |
| 39 | Negative | Negative | Negative |
| 40 | Negative* | Negative | Negative |
| 41 | Negative | Negative | Negative |
| 42 | Negative | Negative | Negative |
| 43 | Negative | Negative | Negative |
| 44 | Negative | Negative | Negative |
| 45 | Negative | Negative | Negative |
| 46 | Negative | Negative | Negative |

Altogether, we tested a total of 196 individual RT-LAMP reactions, comprising 58 spiked RNA and 138 animal RNA samples. We compared the RT-LAMP results to the results of the gold standard RT-qPCR assay, and calculated the sensitivity, specificity, positive and negative likelihood ratio, and positive and negative predictive values. Given that amplification of low abundance targets close to the limit of detection are probabilistic events, multiple replicates are required to estimate the detection probability (or hit rate) at a given concentration. To account for this, every replicate of a spiked and diagnostic sample was treated as an independent observation for statistical analysis, and each result was individually compared to RT-qPCR. A total of 54 of the reactions were positive both by RT-qPCR and RT-LAMP (true positive), 129 were negative by both RT-qPCR and RT-LAMP (true negative), 12 were positive by RT-qPCR and negative by RT-LAMP (false negative) and 1 was considered positive by RT-LAMP while it was negative by RT-qPCR (false positive) (Table S2). The calculated specificity of the test was 99.23%, the sensitivity of the test was 81.82% and the overall accuracy was 93.37%. The sensitivity of the test for samples with a Cq < 35 was 85.7% and with a Cq < 30 was 95.6% (Figure 4B and Tables S3).

## Discussion

We have developed a colorimetric RT-LAMP assay capable of detecting IDV RNA in both experimentally spiked and field-collected samples. The assay demonstrated a high analytical specificity of 99.23% as well as analytical sensitivity of 167 viral RNA copies per µL of target input within a 60-minute reaction time, utilizing a one-step assay and a simple colorimetric detection system for result interpretation. The single tube format allows for simpler handling as fewer manipulations are required, decreasing time from collection to final result. Active viral infections typically produce millions of viral copies, therefore the detection limit in the hundreds of copies range (equivalent to Cq values >30 in RT-qPCR assays) will enable identification of infections during early-stage viral replication and throughout extended viral shedding periods. While sensitivity remains lower than RT-qPCR methodologies, the assay exceeds the analytical performance of most rapid antigen tests and could provide direct evidence of active infection compared to serological approaches.

Primer design incorporated strategically positioned degenerate nucleotides to ensure comprehensive viral lineage coverage, achieving detection capability for at least 90% of published IDV sequences, including the recently identified D/swine/England/126471/2023 lineage from UK swine populations (20). The degenerate nucleotides were specifically integrated avoiding the ends of the primers to minimize potential amplification interference while maximizing lineage inclusivity.

Current IDV detection relies mainly on RT-qPCR and antibody detection, both presenting distinct analytical trade-offs that impact surveillance implementation strategies. RT-qPCR is the gold-standard method and offers superior analytical sensitivity with viral load quantification, enabling precise molecular characterization essential for genetic surveillance and outbreak investigation. However, this approach requires sophisticated laboratory infrastructure, specialized personnel, and complex sample preparation protocols. This limits point-of-care applications and increases costs per sample. Conversely, antibody sampling provides simplified collection protocols with enhanced sample stability and cost-effective population seroprevalence assessment, facilitating large-scale epidemiological studies. Nevertheless, serological methods exhibit inherent limitations due to seroconversion kinetics, cannot distinguish active from historical infections, and may demonstrate cross-reactivity with related viruses, precluding their utility for acute infection diagnosis and real-time outbreak response. These complementary yet functionally distinct methodological constraints necessitate the development of alternative diagnostic platforms that bridge the gap between laboratory-based precision and field-deployable practicality.

RT-LAMP presents a promising point-of-care alternative to RT-qPCR, as its simplified equipment requirements and minimal reaction components enable field-deployable testing with reduced sample processing demands. The colorimetric RT-LAMP approach provides qualitative detection results, facilitating rapid screening of larger sample volumes at reduced per-reaction costs compared to conventional RT-qPCR methodologies. Even though RT-LAMP may be used to differentiate between known lineages, RT-LAMP cannot fully substitute RT-qPCR or sequencing for comprehensive genetic surveillance applications, as its amplification products are difficult to sequence. This is particularly problematic when detecting novel or uncharacterized viral lineages that have not been sequenced and deposited in public databases. Despite these genetic surveillance limitations, RT-LAMP offers significant potential for cost-effective sample triage protocols, enabling preliminary screening before confirmatory molecular characterization through more sophisticated diagnostic platforms.

In summary, our RT-LAMP assay for IDV demonstrated high specificity and sensitivity, indicating its utility as a rapid and accessible diagnostic tool. To further enhance its applicability, future optimisations could focus on integrating the assay into lab-on-a-chip microfluidic platforms or coupling the reaction with a compact detector, allowing for fully automated, portable testing. Further field validation across varied sample types and conditions will be essential, however embedding this RT-LAMP assay into existing surveillance infrastructures can bolster real-time IDV detection and strengthen One Health strategies for pandemic preparedness.

## Acknowledgements

The Isolate D/bovine/France/5920/2014 was kindly provided by Dr Mariette Ducatez, Ecole Vétérinaire de Toulouse (ENVT). This work was supported by a Royal Society Dorothy Hodgkin Research Fellowship [DKR00620 to N.R.] and an Institute for Global Pandemic Planning (IGPP) funded PhD at the University of Warwick, UK [to C.A.dS]. Influenza research at APHA is supported by DEFRA and the devolved Scottish and Welsh Governments previously through FluFutures2 (SE2213) and currently through FluFocus (SE2227). Surveillance sample submissions to APHA were obtained under the National DEFRA-funded surveillance programs SV3041 (swine influenza) and ED1000 and ED200 (bovine respiratory virus).

## References

1. Ng TFF, Kondov NO, Deng X, Van Eenennaam A, Neibergs HL, Delwart E. A Metagenomics and Case-Control Study To Identify Viruses Associated with Bovine Respiratory Disease. J Virol. 2015 May 15;89(10):5340–9.

2. Robinson E, Schulein C, Jacobson BT, Jones K, Sago J, Huber VC, et al. Pathophysiology of Influenza D Virus Infection in Specific-Pathogen-Free Lambs with or without Prior Mycoplasma ovipneumoniae Exposure. Viruses. 2022 Jul 1;14(7).

3. Nedland H, Wollman J, Sreenivasan C, Quast M, Singrey A, Fawcett L, et al. Serological evidence for the co-circulation of two lineages of influenza D viruses in equine populations of the Midwest United States. Zoonoses Public Health. 2018 Feb 1;65(1):e148–54.

4. Gorin S, Fablet C, Quéguiner S, Barbier N, Paboeuf F, Hervé S, et al. Assessment of influenza D virus in domestic pigs and wild boars in France: Apparent limited spread within swine populations despite serological evidence of breeding sow exposure. Viruses. 2019 Dec 24;12(1).

5. Salem E, Cook EAJ, Lbacha HA, Oliva J, Awoume F, Aplogan GL, et al. Serologic evidence for influenza c and d virus among ruminants and Camelids, Africa, 1991-2015. Emerg Infect Dis. 2017 Sep 1;23(9):1556–9.

6. Sreenivasan C, Thomas M, Sheng Z, Hause BM, Collin EA, Knudsen DEB, et al. Replication and Transmission of the Novel Bovine Influenza D Virus in a Guinea Pig Model. J Virol. 2015 Dec;89(23):11990–2001.

7. Song H, Qi J, Khedri Z, Diaz S, Yu H, Chen X, et al. An Open Receptor-Binding Cavity of Hemagglutinin-Esterase-Fusion Glycoprotein from Newly-Identified Influenza D Virus: Basis for Its Broad Cell Tropism. PLoS Pathog. 2016;12(1).

8. Nemanichvili N, Berends AJ, Wubbolts RW, Gröne A, Rijks JM, de Vries RP, et al. Tissue microarrays to visualize influenza d attachment to host receptors in the respiratory tract of farm animals. Viruses. 2021 Apr 1;13(4).

9. Holwerda M, Kelly J, Laloli L, Stürmer I, Portmann J, Stalder H, et al. Determining the replication kinetics and cellular tropism of influenza D virus on primary well-diferentiated human airway epithelial cells. Viruses. 2019 Apr 1;11(4).

10. Borkenhagen LK, Mallinson KA, Tsao RW, Ha SJ, Lim WH, Toh TH, et al. Surveillance for respiratory and diarrheal pathogens at the human-pig interface in Sarawak, Malaysia. PLoS One. 2018 Jul 1;13(7).

11. Hause BM, Ducatez M, Collin EA, Ran Z, Liu R, Sheng Z, et al. Isolation of a Novel Swine Influenza Virus from Oklahoma in 2011 Which Is Distantly Related to Human Influenza C Viruses. PLoS Pathog. 2013 Feb;9(2).

12. Trombetta CM, Marchi S, Manini I, Kistner O, Li F, Piu P, et al. Influenza D virus: Serological evidence in the Italian population from 2005 to 2017. Viruses. 2019 Dec 27;12(1).

13. White SK, Ma W, McDaniel CJ, Gray GC, Lednicky JA. Serologic evidence of exposure to influenza D virus among persons with occupational contact with cattle. Journal of Clinical Virology. 2016 Aug 1;81:31–3.

14. Ferguson L, Eckard L, Epperson WB, Long LP, Smith D, Huston C, et al. Influenza D virus infection in Mississippi beef cattle. Virology. 2015 Dec 1;486:28–34.

15. Alvarez I, Carrera M, Chiapponi C, Ducatez M, El Agrebi N, Faccini S, et al. Developing an integrated approach to assess the emergence threat associated with influenza D viruses’ circulating in Europe. EFSA Supporting Publications. 2024 Feb 6;21(2).

16. Baba MM, Bitew M, Fokam J, Lelo EA, Ahidjo A, Asmamaw K, et al. Diagnostic performance of a colorimetric RT -LAMP for the identification of SARS-CoV-2: A multicenter prospective clinical evaluation in sub-Saharan Africa. EClinicalMedicine. 2021 Oct 1;40.

17. Choi G, Moehling TJ, Meagher RJ. Advances in RT-LAMP for COVID-19 testing and diagnosis. Vol. 23, Expert Review of Molecular Diagnostics. Taylor and Francis Ltd.; 2023. p. 9–28.

18. Ducatez MF, Pelletier C, Meyer G. Influenza D Virus in Cattle, France, 2011–2014. Emerg Infect Dis [Internet]. 2015 Feb 4;21(2). Available from: http://wwwnc.cdc.gov/eid/article/21/2/14-1449_article.htm

19. Salem E, Hägglund S, Cassard H, Corre T, Näslund K, Foret C, et al. Pathogenesis, Host Innate Immune Response, and Aerosol Transmission of Influenza D Virus in Cattle. J Virol. 2019 Apr;93(7).

20. van Diemen PM, Ramsay AR, Bernard M, Floyd T, Byrne AMP, Khatri M, et al. Influenza D virus in Great Britain [Internet]. 2023 [cited 2026 Jan 8]. Available from: https://www.researchgate.net/publication/388946074_Pig_Influenza_D_virus_in_Great_Britain

21. Faccini S, De Mattia A, Chiapponi C, Barbieri I, Boniotti MB, Rosignoli C, et al. Development and evaluation of a new Real-Time RT-PCR assay for detection of proposed influenza D virus. J Virol Methods. 2017 May 1;243:31–4.

22. Lamas A, Azinheiro S, Roumani F, Prado M, Garrido-Maestu A. Evaluation of the efect of outer primer structure, and inner primer linker sequences, in the performance of Loop-mediated isothermal amplification. Talanta. 2023 Aug 1;260.

