## Supplementary material for "Isothermal Detection of Influenza D using RT-LAMP"

**Table S1. RT-LAMP primers for the detection of Influenza D.** Degenerate nucleotides are shown in bold. R: A or G; Y: C or T.

| Primer Name | Sequence (5′ – 3′) |
|---|---|
| IDV_set1.F3 | TGCCTTGTTGAGTACAAGAG |
| IDV_set1.B3 | ATAGGTCCATTATGGATGCA |
| IDV_set1.FIP | ACTCCCTTGTAAACCATAGAAGATGTTTTTTTGCTCATATTGGACAATTCGG |
| IDV_set1.BIP | CGATAGACATAGTGGAGAGCAAAGTTTTTTCCATTGCTCCTCTCTCTGT |
| IDV_set1.LF | TCCATCTATTCCCATTCCTT |
| IDV_set1.LB | AGAAGCACAGACTGTTT |
| IDV_set2.F3 | GGGAATAGATGGAAGTAGCT |
| IDV_set2.B3 | TTTTCATCTTTGAGAACAGCT |
| IDV_set2.FIP | GTG**Y**TTCTCTTTGCTCTCCACTATTTTTTTTGGTTTACAAGGGAGTCAT |
| IDV_set2.BIP | AGGAGCAATGGTTGCATCCATTTTTTTGGTT**R**AAAGTCACATC**R**TTAA |
| IDV_set2.LF | GTCTAT**Y**GGTGTCTTCGACAA |
| IDV_set2.LB | TGGACCTATCAGAGGATAATAGAGA |

**Table S2. Total number of reactions included in statistical analysis.** Only the reactions with spiked and animal collected RNA were considered for these calculations. True positive: Positive both by RT-LAMP and RT-qPCR. True Negative: Negative both by RT-LAMP and RT-qPCR. False Negative: Negative by RT-LAMP and positive by RT-qPCR. False Positive: Positive by RT-LAMP and negative by RT-qPCR.

| | RT-qPCR | | | | |
|---|---|---|---|---|---|
| | Positive | n | Negative | n | |
| RT-LAMP | | | | | Total |
| Positive | True Positive | 54 | False Positive | 1 | 55 |
| Negative | False Negative | 12 | True Negative | 129 | 141 |
| Total | | 66 | | 130 | 196 |

**Table S3. Complete statistical analysis results of the data presented in Table S2**. The sensitivity is the probability the test will be positive when the viral RNA is present. The specificity is the probability the test will be negative when the viral RNA is not present. The positive predictive value is the probability the viral RNA is present if the test is positive, and the negative predictive value is the probability the viral RNA is not present if the test is negative. The accuracy of the test is defined as the overall probability samples are correctly classified. *The disease prevalence refers only to the sample size and not the actual prevalence of the disease.

| **Statistic** | **Value** | **95% CI** |
|---|---|---|
| Sensitivity | 81.82% | 70.39% to 90.24% |
| Specificity | 99.23% | 95.79% to 99.98% |
| Disease prevalence * | 33.67% | 27.10% to 40.75% |
| Positive Predictive Value | 98.18% | 88.42% to 99.74% |
| Negative Predictive Value | 91.49% | 86.56% to 94.72% |
| Accuracy | 93.37% | 88.93% to 96.42% |
